## Supplemental material for "Dynamic spatial coding in parietal cortex mediates tactile-motor transformation"

### Supplementary Information

**Univariate fMRI analyses.** We conducted additional univariate analyses to test whether there were differences in the overall level of BOLD activation between our main conditions of interest, left vs right anatomical coding of tactile stimuli, left vs right external coding of tactile stimuli, left vs right movement goal location and pro- vs anti-pointing task rule. In addition to the preprocessing steps conducted for the multi-voxel pattern classification analysis, T1 images were segmented and the resulting transformation parameters were used to spatially normalize EPI and T1 images to MNI space. EPI images were then smoothed with a 6 mm Gaussian kernel. Participant-level GLMs included the same predictors as the GLM optimized for the multivariate analysis.

We did not identify any regions that showed differences in univariate activation to tactile stimulation on the left vs right foot, to tactile stimulation on the left vs right side of external space or between pro- and anti-pointing conditions. We identified regions that showed significantly higher activation during movement planning to targets on the right as compared to targets on the left (Fig. S3). These regions included a small cluster in the bilateral SMA, the left lateral and medial anterior SPL, M1 and the PMd.

**MVPA analyses with motion parameters included as nuisance regressors.** Our fMRI analyses did not include motion parameters as nuisance regressors in participant GLMs as our unwarping and alignment pre-processing procedure included correction of the fMRI images for movement-related signal intensity changes. To ensure that our results were not affected by any residual movement-related signal changes, we re-ran separate instances of our MVPA analyses that included motion parameters as nuisance regressors in participant GLMs. Results are shown in Fig. S4. In all analyses, locations of clusters are consistent with the analyses without motion parameters. This result suggests that the unwarping procedure had indeed been effective in removing movement-related artifacts.

**MVPA analyses in which trials with missing hand movement data were removed.** We included trials where an interpretable movement trace could not be extracted from the hand videos in our MVPA analyses as participants showed high performance accuracy for trials in which hand movements could be assessed. To ensure that potential errors in trials with missing hand movement data did not affect our MVPA results, we re-ran separate instances of our MVPA analyses excluding these trials. Note that this strategy results in fewer training data for the MVPA classifiers, especially as some runs of data had to be completely excluded. Results of the new analyses are shown in Fig. S5, and show clusters at thresholds both corrected for multiple comparisons ($p < 0.05$ FWE corrected) and uncorrected for multiple comparisons ($p < 0.01$). At the uncorrected threshold, searchlight clusters are located in the same regions as for the MVPA analyses in the main paper, in which we had included trials with missing hand movement data. Thus, results are similar regardless of whether these trials are included or excluded, but power is reduced, as expected, due to the reduction in the total number of trials available for MVPA classification analyses (34% of the total number of trials removed).

##### fMRI analyses with condition duration modelled as 1 TR rather than the full delay interval.

Our GLMs modelled the duration of the sensory processing interval or movement planning interval for each trial (see Fig. 1), as is common in delayed-movement paradigms[1–5]. This strategy aims at modelling sustained neural responses related to sensory processing and movement planning. However, it is also possible that some regions may show only a shorter, more phasic, response following the sensory stimulus or task rule cue. To investigate this possibility, we re-ran separate instances of our MVPA analyses in which the duration of each modelled condition was limited to 1 TR, regardless of the actual length (1-4 TR, see Methods of main paper) of the sensory processing or movement planning interval.

Results of these additional MVPA analyses are shown in Fig. S6. In comparison to results where the full sensory processing or movement planning interval was modelled, decoding of anatomical touch location was found in similar regions, with slightly more lateral clusters in the SPL and more extensive clusters in the insular cortex. External touch location could be decoded bilaterally from regions in the mIPS. Movement goal location could be decoded from a similar fronto-parietal network as in our original analyses, with less extensive clusters in the right hemisphere. Task rule could be decoded from the PPC, as well as additionally from lateral occipital cortex and superior frontal cortex. Note that some additional decoding of task rule could be due to decoding visual differences in the task rule cue or transient coding of the task rule[6]. Such responses are likely prominent during initial decoding but no longer detectable when a longer time interval is analyzed.

Univariate fMRI analyses (Fig. S7) showed higher responses during movement planning to the right versus left side of space in the left SMA, anterior SPL, M1 and the PMd, consistent with the results we obtained in the main paper, when we modelled the full sensory processing or movement planning interval. No regions showed higher responses during movement planning to the left versus right side of space. A small cluster in the left insula showed higher responses to sensory stimuli on the right foot compared to the left foot. Clusters in the left somatosensory and superior temporal cortex showed higher responses to touch on the left vs right side of space. Lastly, clusters in the left posterior PPC, and left superior frontal cortex showed higher responses to the anti-movement task compared to the pro-movement task. None of the other contrasts (left vs right anatomical touch location, right vs left external touch location, pro vs anti task rule) showed any clusters with statistically significant differences in responses.

#### Supplementary Figures

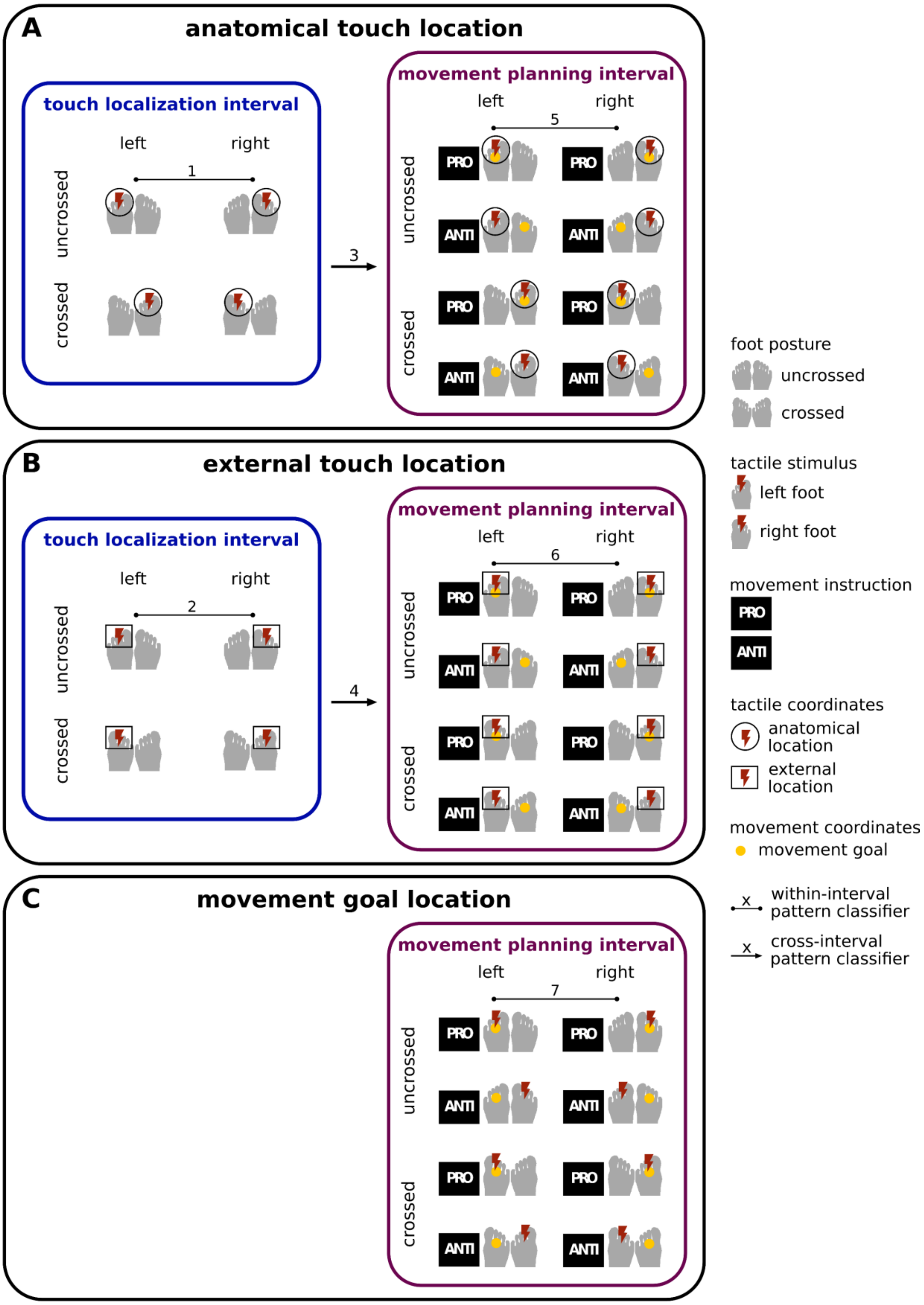


**Fig. S1** Full design scheme of MVPA decoding analyses. Within-interval pattern classification refers to analyses in which both training and test data stemmed from the same trial interval, which was either fMRI data from the touch localization interval (blue boxes, left) or the movement planning interval (purple boxes, right). Cross-interval classification refers to analyses in which training data from the touch localization interval were used to predict the classification labels of test data from the movement planning interval. **A** Pooling of conditions to decode anatomical touch location (left vs. right foot). **B** Pooling of conditions to decode external touch location, i.e., spatial side of touch. Note, that foot crossing dissociates anatomical from external location (denoted by the lightning-bolt in circled vs. squared boxes on the right side of the figure). **C** Pooling of conditions to decode the location of the movement goal. The combination of foot posture, stimulation side, and pro/anti-pointing dissociates sensory and movement coding.

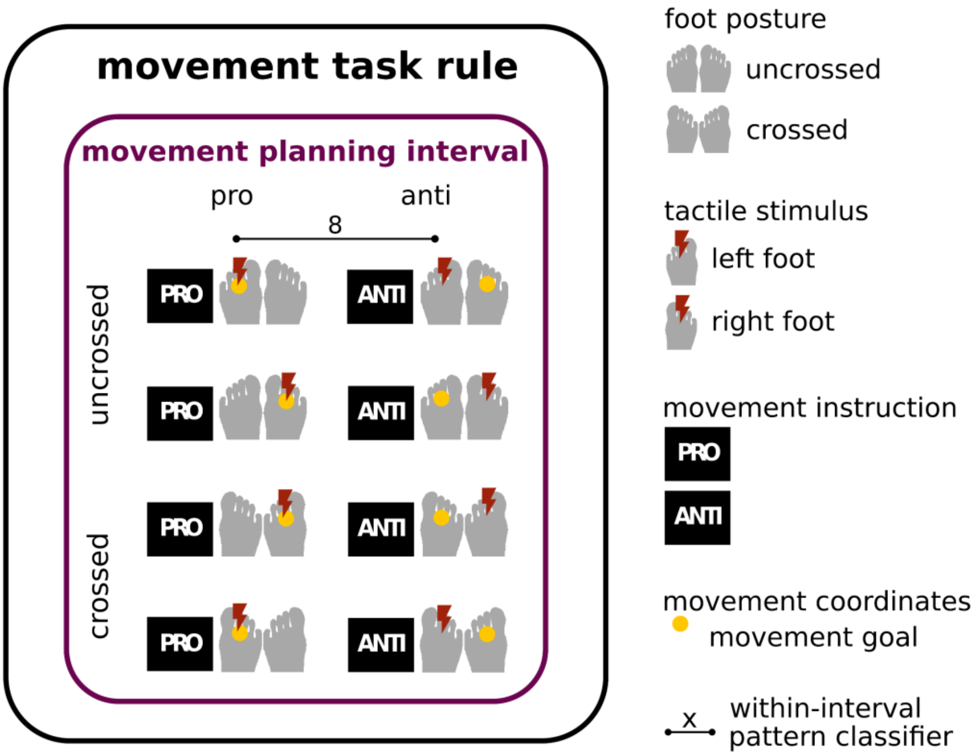


**Figure S2.** Full design scheme of the task rule decoding analysis to distinguish patterns of responses evoked by pro- vs. anti-pointing movements during the movement planning interval.

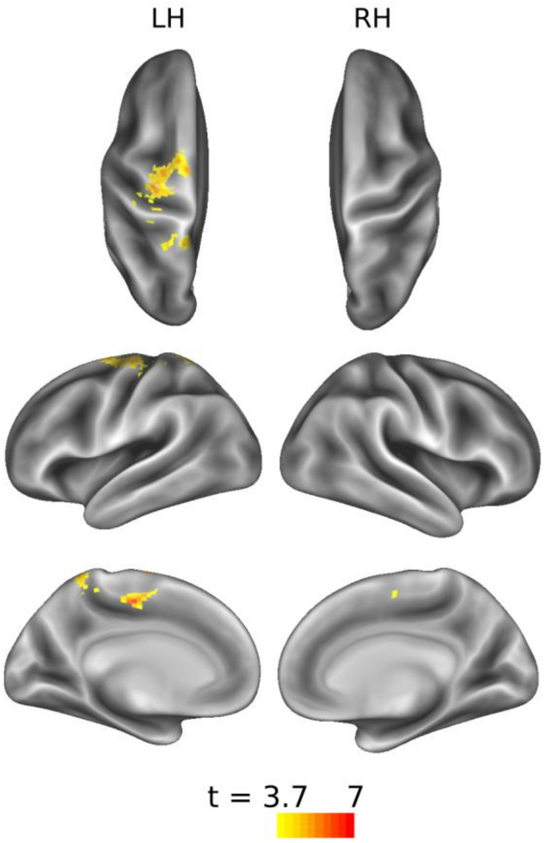


**Figure S3.** Regions showing higher BOLD activation during movement planning to targets on the right as compared to targets on the left. Results show clusters with significantly higher group-level activation for right vs left movement planning, corrected for multiple comparisons using a cluster-based permutation test (FWE, p<.05). LH, left hemisphere; RH, right hemisphere.

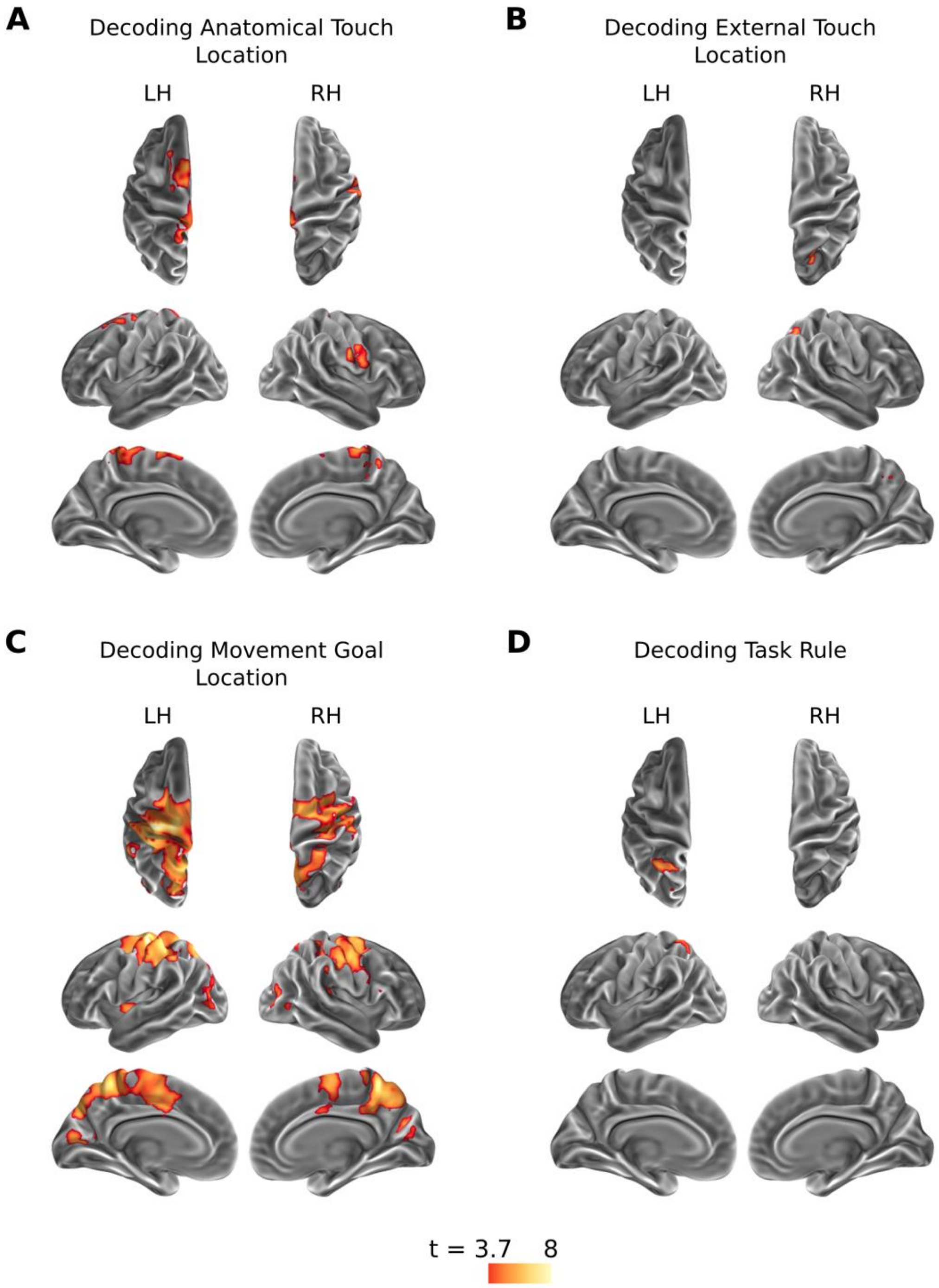


**Figure S4.** Results of decoding analyses with motion parameters included as nuisance regressors in participant GLMs. **A** MVPA results for decoding anatomical touch location (right vs. left foot) during the touch localization interval. **B** MVPA results for decoding external touch location (right vs. left side of space) during the touch localization interval. **C** MVPA results for decoding movement goal location (pointing movement to the foot on the right or left side of space) during the movement planning interval. **D** MVPA results for decoding task rule (pro- vs. anti-pointing movement) during the movement planning interval. Results show clusters with significantly higher group-level activation, corrected for multiple comparisons using a cluster-based permutation test (FWE, $p<.05$). LH, left hemisphere; RH, right hemisphere.

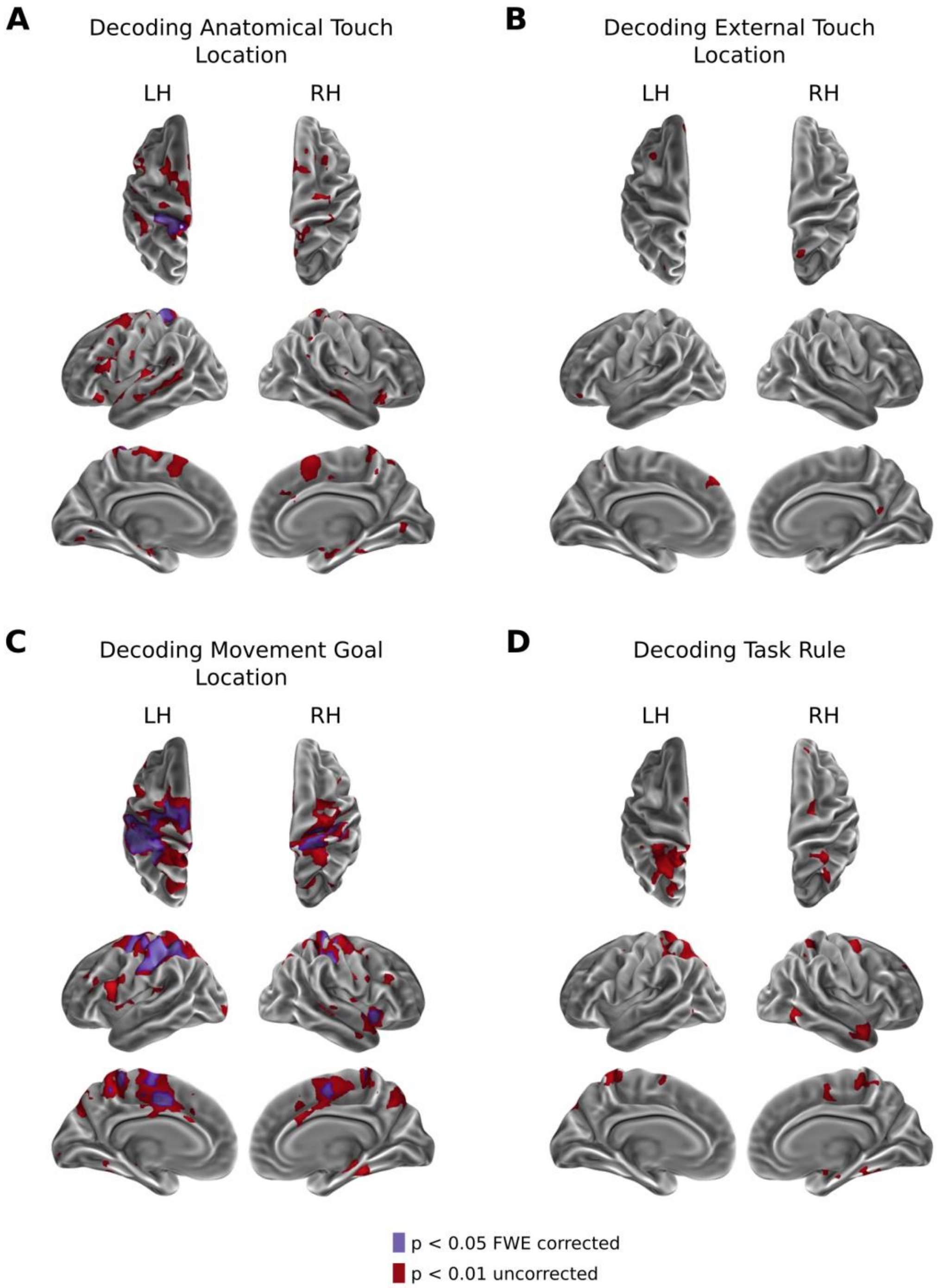


**Figure S5.** Results of MVPA decoding analyses in which we removed all trials in which an interpretable movement trace could not be extracted from the hand videos. **A** MVPA results for decoding anatomical touch location (right vs. left foot) during the touch localization interval. **B** MVPA results for decoding external touch location (right vs. left side of space) during the touch localization interval. **C** MVPA results for decoding movement goal location (pointing movement to the foot on the right or left side of space) during the movement planning interval. **D** MVPA results for decoding task rule (pro- vs. anti-pointing movement) during the movement planning interval. Results corrected for multiple comparisons using a cluster-based permutation test (FWE, $p<.05$) are shown in purple, and results at a threshold uncorrected for multiple comparisons ($p < 0.01$) are shown in red. LH, left hemisphere; RH, right hemisphere.

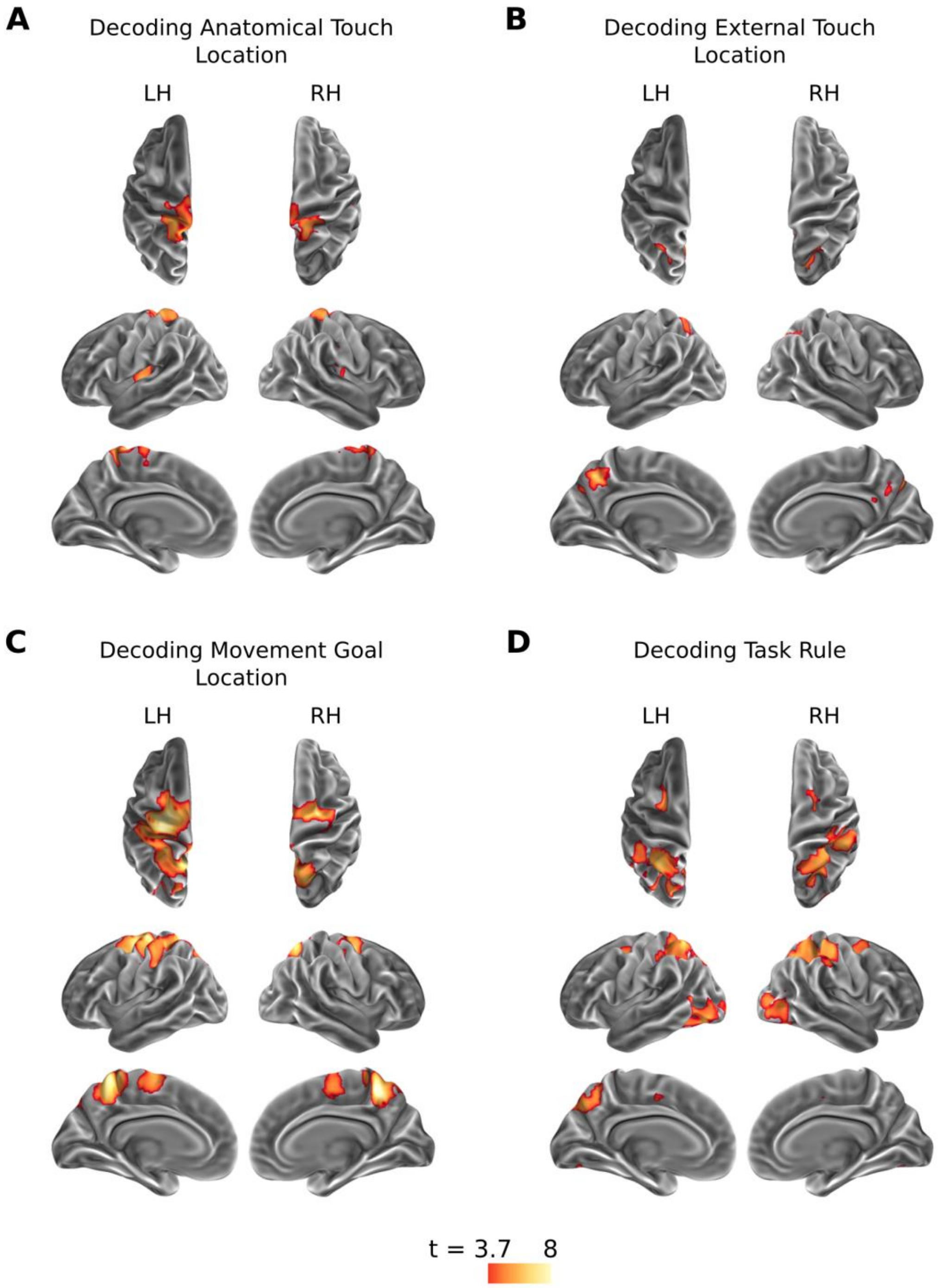


**Figure S6.** Results of MVPA decoding analyses where the modelled duration was reduced to 1 TR for all trials independent of the actual duration of the touch localization or movement planning interval. **A** MVPA results for decoding anatomical touch location (right vs. left foot) during the touch localization interval. **B** MVPA results for decoding external touch location (right vs. left side of space) during the touch localization interval. **C** MVPA results for decoding movement goal location (pointing movement to the foot on the right or left side of space) during the movement planning interval. **D** MVPA results for decoding task rule (pro- vs. anti-pointing movement) during the movement planning interval. Results show clusters with significantly higher group-level activation, corrected for multiple comparisons using a cluster-based permutation test (FWE, $p<.05$). LH, left hemisphere; RH, right hemisphere.

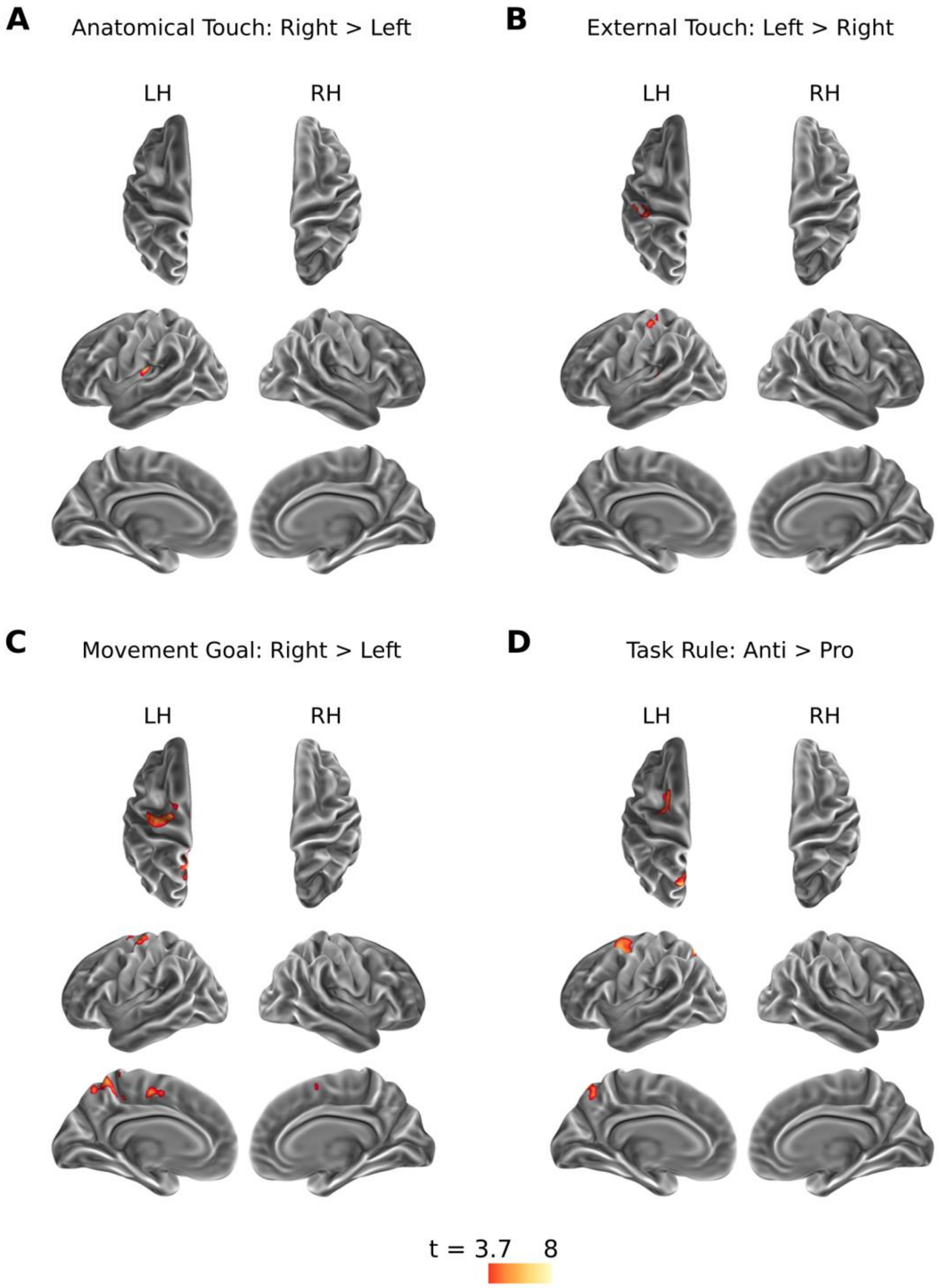


**Figure S7.** Results of univariate analyses where the modelled duration was reduced to 1 TR for all trials independent of the actual duration of the touch localization or movement planning interval. **A** Regions showing higher BOLD activation during sensory processing of stimuli with anatomical location on the right foot as compared to the left foot. **B** Regions showing higher BOLD activation during sensory processing of stimuli with external location on the left side of space as compared to the right side of space. **C** Regions showing higher BOLD activation during movement planning to targets on the right as compared to targets on the left. **D** Regions showing higher BOLD activation during anti-movement planning as compared to pro-movement planning. Results show clusters with significantly higher group-level activation, corrected for multiple comparisons using a cluster-based permutation test (FWE, p<.05). LH, left hemisphere; RH, right hemisphere.
